# JavaDL: a Java-based Deep Learning Tool to Predict Drug Responses

**DOI:** 10.1101/2020.05.04.077701

**Authors:** Beibei Huang, Lon W. R. Fong, Rajan Chaudhari, Zhi Tan, Shuxing Zhang

**Author notes:** Corresponding authors: Shuxing Zhang.

## Abstract

**Motivation:** Accurate prediction of drug response in each patient is the holy grail in personalized medicine. Recently, deep learning techniques have been witnessed with revival in a variety of areas such as image processing and genomic data analysis, and they will be useful for the coming age of big data analysis in pharmaceutical research and chemogenomic applications. This provides us an impetus to develop a novel deep learning platform to accurately and reliably predict the response of cancer to different drug treatments.

**Results:** In this study, we describe a Java-based implementation of deep neural network (DNN) method, termed JavaDL, to predict cancer responses to drugs solely based on their chemical features. To this end, we devised a novel cost function by adding a regularization term which suppresses overfitting. We also adopted an “early stopping” strategy to further reduce overfit and improve the accuracy and robustness of our models. Currently the software has been integrated with a genetic algorithm-based variable selection approach and implemented as part of our JavaDL package. To evaluate our program, we compared it with several machine learning programs including SVM and kNN. We observed that JavaDL either significantly outperforms other methods in model building and prediction or obtains better results in handling big data analysis. Finally, JavaDL was employed to predict drug responses of several highly aggressive triple-negative breast cancer cell lines, and the results showed robust and accurate predictions with r^2^ as high as 0.80.

**Availability:** The program is freely available at https://imdlab.mdanderson.org/JavaDL/JavaDL.php.

## 1. Introduction

Over the past ten years, many machine-learning methods have been developed to tackle the problem in biomedical research [1, 2]. Recently a new branch of machine learning known as deep learning has been gaining significant attention [3-8]. In particular, the resurgence of neural networks took place around 2005 when more efficient training algorithms were developed and improvements in overfitting were made. This has led to the success of deep neural networks with many applications to protein structure prediction and drug development [9-13]. Moreover, deep learning has been employed to diagnose diseases based on medical images [14-16].

In the canonical configuration, a normal deep neural network (DNN) consists of an input layer where an input signal is fed, an output layer where predictions are generated, and several hidden (middle) layers which capture features during the training[17]. DNN is also considered a type of representation learning, in that it allows a machine to be fed raw data and automatically discover the representations needed for detection. During the training process, the raw data in the form of certain signals are fed into the input layer and then transported from one hidden layer to another, in each one leaving a trail forming a certain increasingly abstractive pattern. For instance, in image processing the raw data are arrays of pixel values, while in quantitative structure-activity relationship (QSAR) modeling it may be descriptors including a variety of physical chemical properties of compounds. Such applications involve a large amount of input data, exposing the drawbacks of previous multilayer neural network (MNN) programs, which only accept limited numbers of input descriptors and have limited numbers of hidden layers and neurons in a hidden layer. Due to these limitations, networks with only a single hidden layer were largely abandoned [18].

One of the major issues with artificial neural networks is that the models are significantly complex, since neural networks usually have large numbers of layers containing many neurons. The number of connections in these models is astronomical, easily reaching the millions, and overfitting thus becomes very common [19]. In general, there is a direct trade-off between overfitting and model complexity. Inadequately complex models may not be powerful enough to capture all the information necessary to solve a problem, but overly complex ones (especially with a limited amount of data) tend to run into the risk of overfitting [19]. Besides overfitting, another challenge in drug response prediction comes from the activity cliff that is formed when a pair of structurally similar molecules display a large difference in potency [20, 21]. From the perspective of chemical structures, the activity cliff indicates a lack of assayed compounds in the surrounding space. This can also lead to poorly generalized models and overfitting. Furthermore, the distance between compounds is defined based on their relationship to neighboring compounds. Hence almost all current published prediction models are flawed to some degree due to the limitations that arise from one or more of the above problems.

In the present study, we aim to develop a novel method and build robust models that accurately predict drug response in cancer cells. To this end, we designed an approach which employs deep learning algorithms for drug activity prediction. Our software package, termed JavaDL, integrates several of the latest improved techniques, including regularization, dropout, and early stopping, to mitigate the issues caused by overfitting and activity cliffs. In order to assess its robustness, we used two very different datasets: a Caco-2 dataset for permeability prediction and a hERG dataset for cardiovascular toxicity prediction. We also evaluated the ability and robustness of JavaDL to perform big data analysis using the Merck Molecular Activity Challenge dataset from Kaggle. Finally, our software was employed to predict the drug response of cancer cells using data we recently curated and experimentally measured. The results show that JavaDL could obtain significantly improved prediction and have broad applicability in a variety of datasets as well as a high capability to handle big data problems.

## 2. Methods

### 2.1 Datasets

For model building and prediction evaluation, we compared JavaDL with several published studies using the exact same datasets. The first dataset was curated by our group and it has been used to build Caco-2 permeability prediction models with kNN and SVM [22, 23]. The whole dataset contains 174 compounds with 334 descriptors calculated using MOE. Another dataset contains hERG blockers with their corresponding known hERG inhibition activity (pIC_50_) obtained from previous publications [24]. This dataset includes 639 hERG active and inactive compounds. Similarly, MOE was used to calculate the molecular descriptors which were normalized to avoid disproportional weighting for both datasets [24]. In addition, to evaluate the ability of JavaDL to handle big data, we obtained a dataset for the Merck Molecular Activity Challenge on Kaggle. The original training data set contained 1,569 compounds with 4,505 descriptors, which after washing shrank to 1,983 descriptors [25]. Finally, we applied JavaDL to predict the triple negative breast cancer (TNBC) cell response to drugs. The experimental data of cancer cell response to drugs were collected from the MIPE project at the National Center for Advancing Translational Science (NCATS) [26] and our own laboratory’s data. We focused on triple-negative breast cancer (TNBC) due to our own research interest, but the approach can be easily applied to other cancer cell lines or even other diseases. In particular, we selected four breast cancer cell lines, HCC-1937, MDA-MB-436, MDA-MB-231, and MDA-MB-453, representing four different TNBC molecular subtypes. The dataset contains 274 compounds used in all four cell lines. Each compound is described by 205 molecular descriptors and four half maximal activity (log) concentration (LAC50) values (one for each cell line). Although internal cross validation is generally considered sufficient to justify model predictive power [27, 28], many researchers have argued that external validation is crucial. In this study compounds in our datasets are divided into training and test sets via a rational approach based on the Sphere Exclusion algorithm [29, 30].

### 2.2 Techniques to address challenges in DNN

#### 2.2.1 Activation function

For our program, we essentially adopted a backpropagation algorithm to train the deep neural network. Based on our experience and other reports [19, 31-35], we included five total layers for our deep neural networks, with three hidden layers. Assuming five nodes in the input layer and one node in the output layer, three hidden layers each containing 20 hidden nodes makes the total number of variables (weight variables) 920. In contrast to frequently used activation functions in previous studies [34], on each hidden node we adopted an activation function 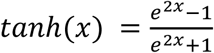 (**Fig. 1)**, with which one of the advantages is to avoid bias in the gradient during training. For the loss function we chose mean squared error. In the backpropagation process we employed the stochastic gradient descent algorithm for optimization. Additionally, JavaDL has been implemented in a way to deal with data featuring different scales and complexity. For instance, to build a model from a limited dataset, the 5-layer structure with each layer containing 20 neurons is sufficient. However, for large datasets such as those in the Merck Molecular Activity Challenge competition, JavaDL can grow accordingly the number of layers and neurons with corresponding increase of the connections between neighboring layers.

**Figure 1.**
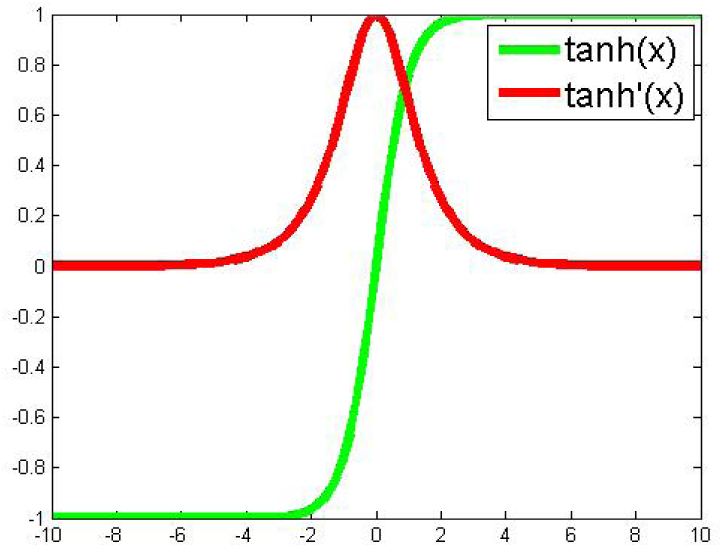
Activation function based on the Tanh function tanh(x) and its derivative tanh’(x)

#### 2.2.2 Regularization

To address the overfitting issue, we adopted a common regularization technique: addition of a regularization term to the cost function. It is based on the relation between the regularized overall squared-error cost function and the correlation coefficient *q*^2^. Given a training set of *m* examples, the overall squared-error cost function *J(W, λ)* is considered below:

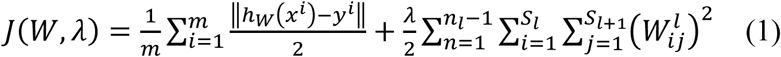

where *n*_*l*_ denotes the number of layers in the network. *S*_*l*_ represents the number of nodes on lay *l*, and 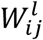 denotes the weight of the connection between node *i* on lay *l* and node *j* on lay *l* + 1. The first term is an average sum-of-squares error. For simplicity, we neglect the bias term, but it can be easily taken into account. The second term is the weight decay term, which suppresses overfitting via tuning the value of λ to decrease the magnitude of the weights. The method of minimizing the cost function by searching for the optimal value of *λ* is introduced in [36]. This method, called Bayesian Regularized Neural Networks, involves incorporating Bayes’ theorem into the regularization scheme. It has been frequently used in current research.

We defined our own cost function by replacing the average sum-of-squares error with *– q*^*2*^

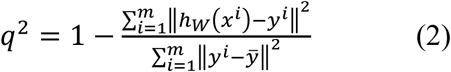

where 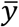 is the average actual activity of the training set. The relation in **Eq. 3** guarantees the consistency between the results from the minimization of *J* and the maximization of *q*^*2*^; therefore we consider only the minimization of cost *J(W, λ)* hereafter.

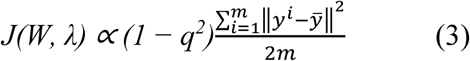

#### 2.2.3 Dropout and early stopping

It has been suggested to use at least two hidden layers with a minimum of 250 neurons in each layer [34]. Since our DNNs have a large number of layers containing many neurons, the increasing number of connections between these neurons makes the corresponding Hessian matrix in the backpropagation process increase complexity by *O(N*^*2*^), where *N* is the layer size, and the number of variables in these models can reach into the millions easily. To improve efficiency, JavaDL adopts a process of randomly “dropping out” some neurons in the hidden layers during the training process, which involves temporarily removing the randomly selected neurons from the network, along with all their incoming and outgoing connections [37]. This strategy reduces the complexity of JavaDL’s backpropagation computation and ameliorates overfitting as well. Several other methods have also been used to implement the early stopping strategy in JavaDL [38][39]. The default early stopping criterion of JavaDL is based on the evaluation of the cost function value on a test set.

### 2.3 Implementation of JavaDL

#### 2.3.1 The overall framework

The overall framework of our system consists of four steps as illustrated in **Fig. 2**. The first step is to curate compound structures and their corresponding bioactivities from literature and other online sources. In the second step we calculate the descriptors of compounds with MOE [40] and CDK[41]. The third step is to clean the data as described below, and in the final step the data is used as input to JavaDL to train the program and build models. We chose XML as the format for parameter input to provide extra flexibility for the use of the program: since it is verbose, self-describing and extendable, it can be easily incorporated in other applications. Our software was developed using DL4J, an open-source, distributed deep-learning library. For easy-to-access and easy-to-use, the software package is made available online along with an instruction manual (http://imdlab.org/JavaDL/) for download upon request.

**Figure 2.**
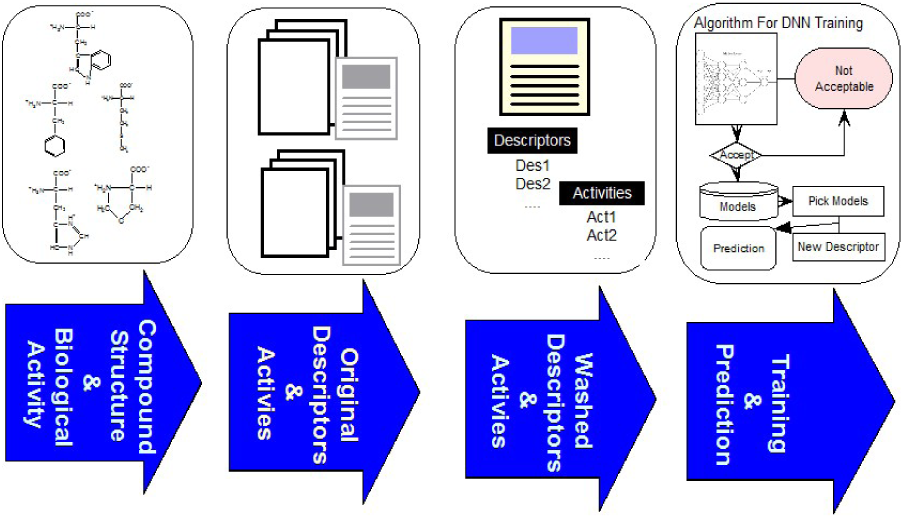
The overall framework of JavaDL including data processing and DNN implementation.

#### 2.3.2 Data cleaning

Online or published data are rich but often “dirty,” and thus generally need further cleaning and curation. Detection of true outliers is of particular importance; therefore, outlier separation from the main data is included in the data-washing step. The premise here is that similar structures have similar biological activity and that the multidimensional response surface to densely sampled data follows a normal distribution [32, 42]. Therefore, we adopted the Pauta criterion to detect abnormal activity-value points, defining a compound with activity value over three standard deviations (3σ) from the mean as an outlier. However, to be statistically significant, the number of compounds similar to the outlier in the descriptor space should be more than 10.

Currently over 5000 molecular descriptors have been reported [43, 44], thus requiring selection of the most relevant and independent descriptors that can represent specific properties of the chemical entities, with smaller numbers of descriptors preferred. Principal components analysis (PCA) has been frequently employed to reduce the dimension of the descriptor matrix[45, 46]. Our initial implementation was based on PCA: given a descriptor matrix 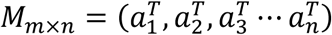, in which *n* denotes the number of compounds and 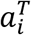 represents the vector containing *m* descriptors; the algorithm is then described in the following steps:

*Step 1*. Calculate the mean of each column 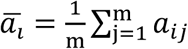 and substract the mean 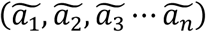;

*Step 2*. Calculate the covariance matrix;

*Step 3*. Calculate the eigenvectors and eigenvalues of the covariance matrix *M*_*cov*_, then order them by eigenvalues from highest to lowest;

*Step 4*. Select *P* important eigenvectors from the highest *P* eigenvalues to compose a feature matrix M_feature,n×p_ = v_1_, v_2_, ⋯ v_p_;

*Step 5*. With the new feature matrix, we derive a new low-dimension matrix via 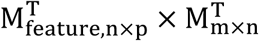 for model building.

However, during the process, the challenge was that the descriptor covariance is too high and *P* is difficult to select. To overcome this problem, we implemented a simple workaround: instead of considering the covariance of descriptors between two compounds, we considered the correlation between two descriptors. In the descriptor matrix *M*_*m*×*n*_ = *(b*_1_, *b*_2_ ⋯ *b*_*n*_*), b*_*i*_ represents the *i*th descriptor vector. We assume if there is a high correlation between descriptor vectors *b*_*i*_ and *b*_*j*_, they are not considered orthogonal to each other, since either alone carries enough information to distinguish signals during the training process. Our final implementation with a simple algorithm to exclude the redundant descriptors is as follows:

1. *For* i *= 1 to* n
2. *For* j *= i to* n
3. *If* corr(b_i_, b_j_) *>* threshold
4. *Delete column* j *from* M
5. *End if*
6. *End for*

where *corr(b*_*i*_, *b*_*j*_) is the function calculating the value of correlation between *b*_*i*_ and *b*_*j*_, and “threshold” denotes the tolerance value for the correlation coefficient. **Eq. 2** is used to evaluate the correlation directly. This is much more efficient and effective.

#### 2.3.3 Deep learning implementation

Our JavaDL employs deep layer neural networks as the primary training engine. A variable selection procedure was also implemented as an option, in particular for small datasets, where for each predefined number of variables it seeks to optimize the models with the highest correlation coefficient (*q*^*2*^) (**Eq. 2**) for both internal training set and external test set. **Fig. 3** shows the flowchart of JavaDL training process, in which the backpropagation algorithm is utilized. The most time-consuming step in this algorithm is the calculation of the stochastic gradient descent (SGD) in each layer and the weight (neuron connections) adjustment to minimize the errors derived from our overall squared-error cost function *J(W,λ)* (**Eq. 1**) in each iteration. In other words, through the backpropagation algorithm, JavaDL knows how to reduce the error from the cost function, rather than from blindly wandering in space or being guided by a scalar quantity such as the random walk in the Metropolis algorithm.

**Figure 3.**
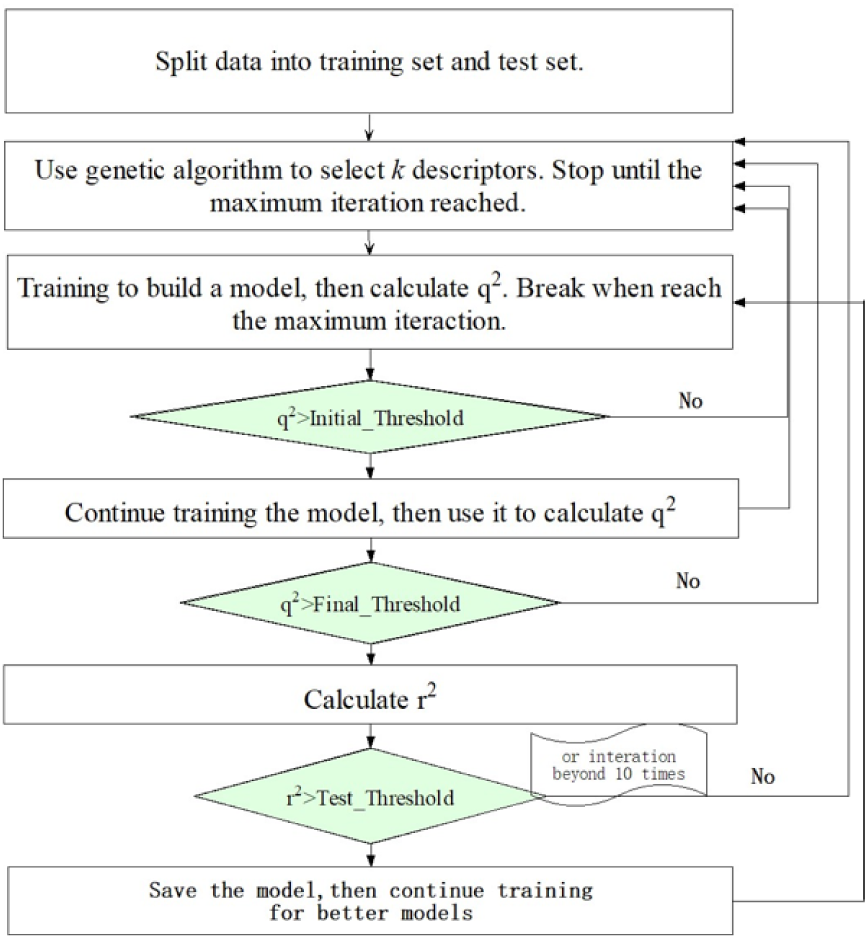
Overall flowchart of DNN Algorithm.

Generally, the time complexity is determined by the number of descriptors chosen and the structure of the neural network. Let *k* denote the number of descriptors picked from *K* total descriptors, *m, n* the number of layers and neurons, respectively, in each layer (assume each layer contains the same number of neurons), and *d* the size of the mini-batch representing the minimum iteration time in the training process, the time complexity is then 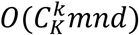.

## 3. Results and Discussion

### 3.1 Comparison with other machine learning programs

Before training, we first performed data wash as described previously and then divided the data into two sets using the Sphere Exclusion (SE) algorithm [29, 30]. For the Caco-2 dataset, one is for training with 80 compounds and the other is for testing with 20 compounds. It should be noted that the division process is executed each time after changing the descriptors during iterations. The same procedure was followed for the hERG dataset, with 133 compounds for training and 14 compounds for testing. Results from JavaDL are compared with those obtained by kNN and SVM methods as shown in **Table 1** and **Table 2**. Also as illustrated in **Fig. 4** our models show high correlation between the actual and predicted activities for the test sets.

**Table 1.**
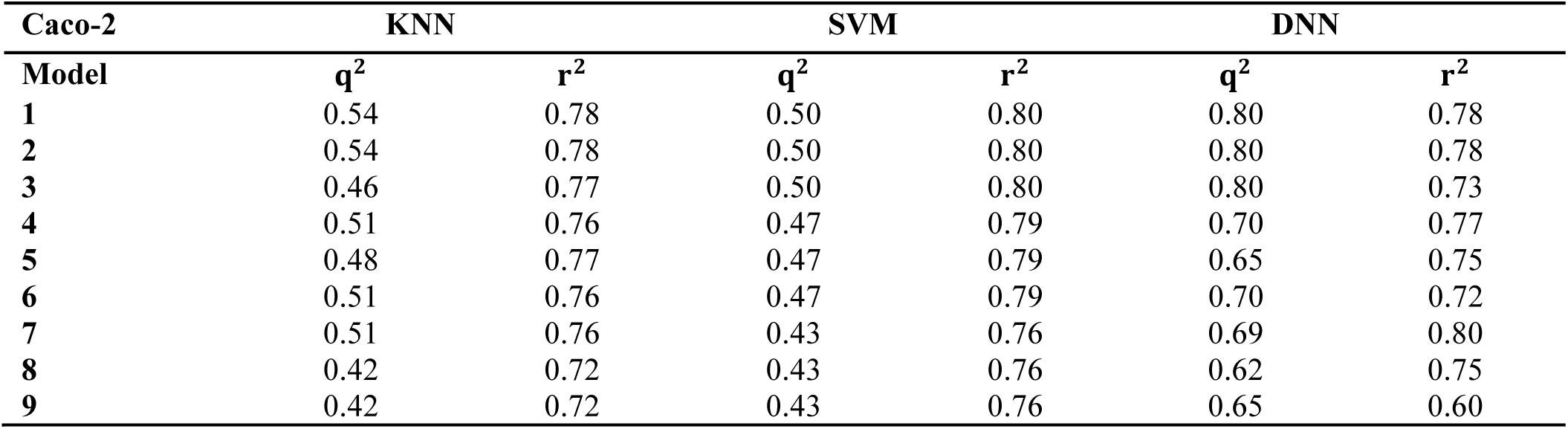
Method Comparison with the Caco-2 Dataset

**Table 2.**
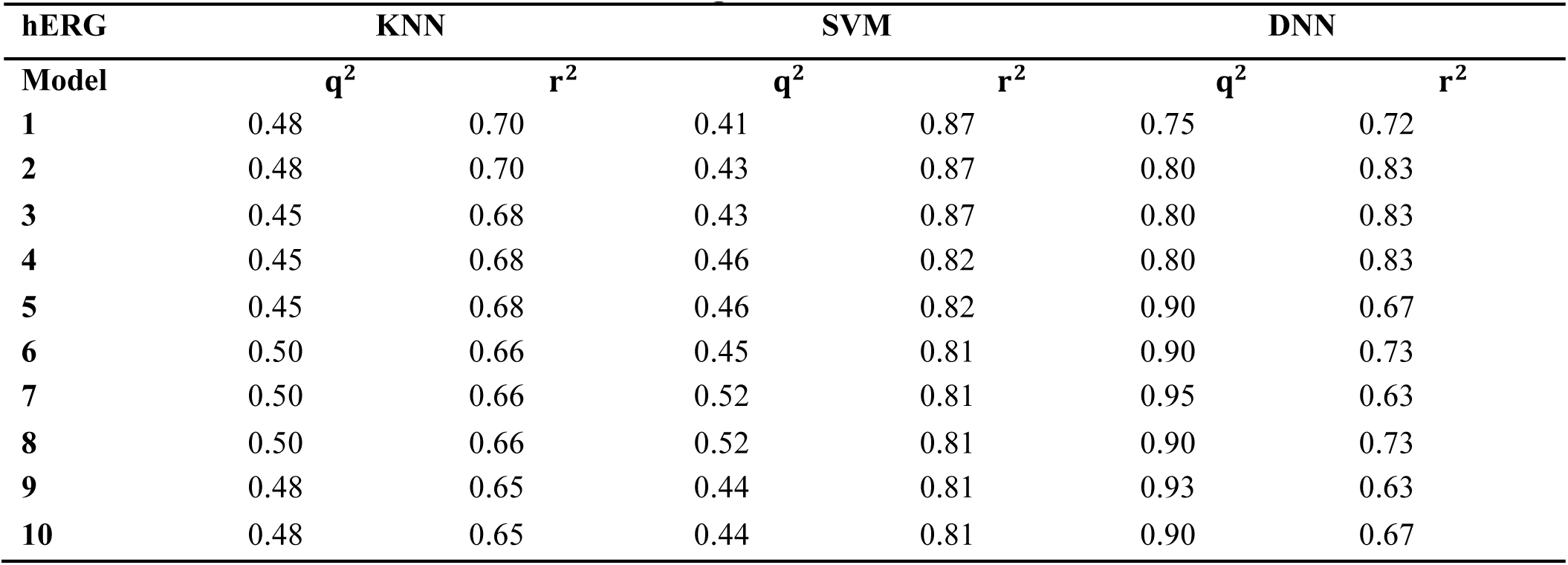
Method Comparison with the hERG Dataset

| hERG | KNN |  | SVM |  | DNN |  |
| --- | --- | --- | --- | --- | --- | --- |
| Model | $q^2$ | $r^2$ | $q^2$ | $r^2$ | $q^2$ | $r^2$ |
| 1 | 0.48 | 0.70 | 0.41 | 0.87 | 0.75 | 0.72 |
| 2 | 0.48 | 0.70 | 0.43 | 0.87 | 0.80 | 0.83 |
| 3 | 0.45 | 0.68 | 0.43 | 0.87 | 0.80 | 0.83 |
| 4 | 0.45 | 0.68 | 0.46 | 0.82 | 0.80 | 0.83 |
| 5 | 0.45 | 0.68 | 0.46 | 0.82 | 0.90 | 0.67 |
| 6 | 0.50 | 0.66 | 0.45 | 0.81 | 0.90 | 0.73 |
| 7 | 0.50 | 0.66 | 0.52 | 0.81 | 0.95 | 0.63 |
| 8 | 0.50 | 0.66 | 0.52 | 0.81 | 0.90 | 0.73 |
| 9 | 0.48 | 0.65 | 0.44 | 0.81 | 0.93 | 0.63 |
| 10 | 0.48 | 0.65 | 0.44 | 0.81 | 0.90 | 0.67 |

**Figure 4.**
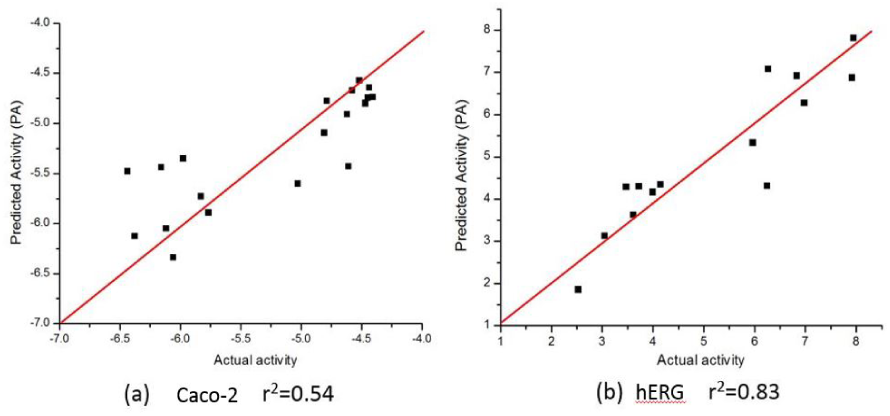
Predicted vs. actual activities obtained for test sets. **(a)** Prediction of Caco-2 permeability using JavaDL with five descriptors: BCUT_ PEOE_3, GCUT_SLOGP_1, mr, a_base, vsa_base. **(b)** Prediction of hERG activity with five descriptors: GCUT_PEOE_3, reactive, SlogP_V SA1, SlogP_V SA9, vdw_area.

The statistical parameters used to assess models, including *q*^*2*^ and *r*^*2*^, demonstrate the comparable performance of JavaDL to other machine learning algorithms, particularly when datasets are small.

We also examined our JavaDL program using a large data set from the Merck Molecular Activity Challenge as described above. Since this would make the training computationally intensive, we have implemented a parallel computing function for JavaDL so that it can efficiently handle big data with a large number of compounds and high dimensional space (number of descriptors). With 250 compounds in the test set, our model achieved robust predictions with *q*^*2*^ = 0.99 and *r*^*2*^ = 0.65. These results demonstrate significant superiority to all published models released by the Merck Molecular Activity Challenge where *r*^*2*^ was below 0.49. Our models and predictions are shown in **Fig. 5**.

**Figure 5.**
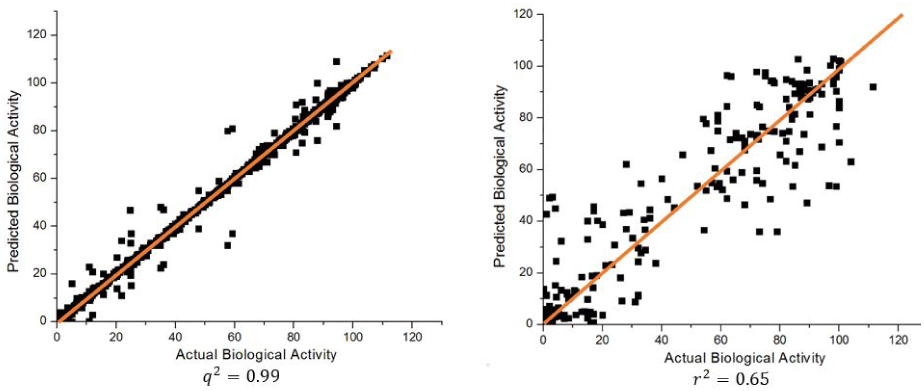
Predicted vs. actual biological activities for the Merck Molecular Activity Challenge big data set in <u>Kaggle</u> competition. The model was built using 1,569 compounds with 4,505 descriptors. Among them 250 compounds are picked out randomly for test predictions, and the best model derived q^2^=0.99, shown in the left panel, and r^2^=0.65, shown in the right panel.

### 3.2 Predicting cancer response to drugs

As discussed, it is critical to predict individual cancer cell response to different drugs. Such information can provide insight into what drugs can be used to treat what type of cancer with the highest sensitivity. With TNBC screening data from NCATS and our own experiments, we performed data pre-processing, and the total number of compounds for each of the four TNBC cell lines was reduced to 186, 163, 146 and 172, respectively. In each group we randomly selected 40 compounds as the external test set, and set 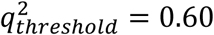 and 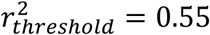 as criteria to determine if a model is acceptable. Since the data is sufficiently large and JavaDL can handle big data efficiently, we took into account all descriptors of the compounds for model building and final predictions. The measured and predicted activities with our best models are shown in **Fig. 6**.

**Figure 6.**
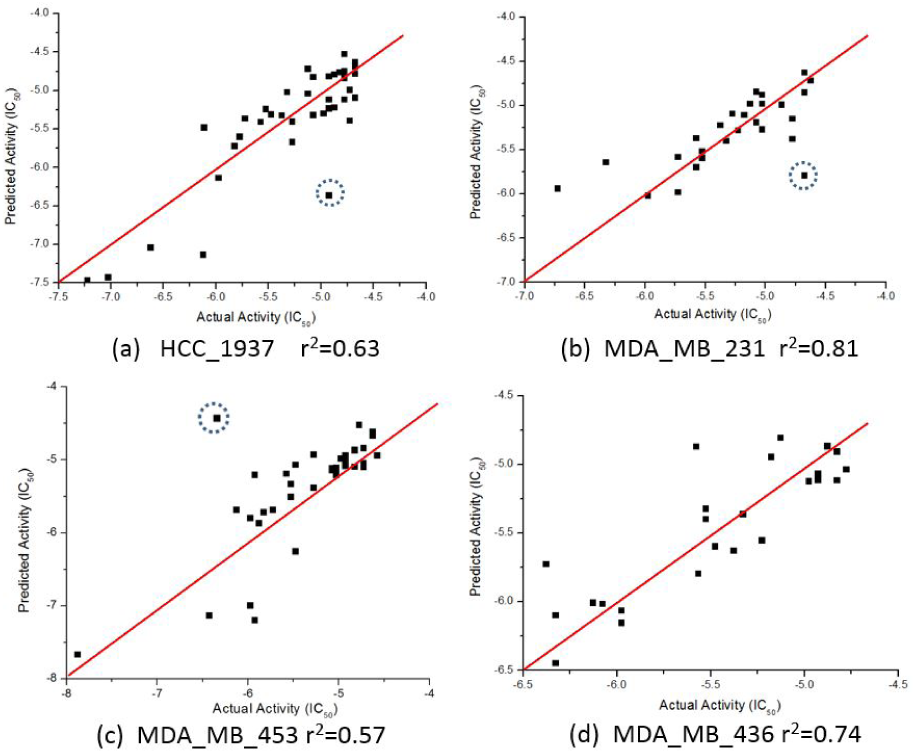
Predicted vs actual activities (LAC50) of drugs in TNBC, including HCC_1937, MDA_MB_436, MDA_MB_231, and MDA_MB_453. The red-circled compound 3-Methyladenine as an outlier was not accurately predicted. Further analysis shows that no compound in the training set is similar enough to 3-Methyladenine while there are always similar compounds to the accurately predicted ones (blue-circled Afatinib and UNC-0638).

### 3.3 Discussion

In this study, we present a new deep-learning method for predicting the efficacy of small-molecule anti-cancer therapeutic agents. The novelty of this implementation lies in multiple aspects. The first is that prediction of drug response has been challenging due to the intrinsic complexity of cancer cells and various unknown cell survival mechanisms [47, 48]; DNN is exactly designed to handle this type of “black-box” problems through a continuous self-learning process. Second, we have implemented a variety of strategies to address several frequent issues including data cleaning, overfitting, outliers, and program complexity. Third, we designed a new scoring activation function and cost function. Fourth, JavaDL is able to predict the response of cancer cells to drugs using only structural information from compounds, and it can be easily expanded to include cell gene profile change upon drug treatment for predictions. Finally, DNN is combined with variable selection in this parallelized implementation and we could identify specific descriptors that are critical for activities. Such information is tremendously helpful to guide actual rational drug design.

Using two very well studied datasets (hERG and Caco-2) as benchmarks, we obtained robust results demonstrating the superiority of our multiple-layer DNN techniques to other traditional machine learning techniques such as kNN and SVM. It also suggests that our models could effectively capture the abstract relation between the structural features of chemical compounds and their activities. Also as shown, JavaDL was successfully employed to predict the response of aggressive TNBC cell lines to different drugs, which has been a challenge in personalized cancer therapy. Moreover, this indicates that JavaDL may potentially be used as a general screening tool to predict activities of novel compounds in different cancer cells, and thus helping to lower costs of anticancer therapeutics screening. It is mostly worth to mention that, as one of the signified features of deep learning, our prediction of the Merck Molecular Activity Challenge dataset demonstrated the capacity of JavaDL to handle big data problems with a large number of points in a high dimensional space.

As mentioned, the variable selection feature in this implementation is particularly useful for rational molecular design, and identification of selected descriptors, when combined with chemical structures, can be employed to interpret the molecular structure-activity relationship and guide lead optimization in drug development. For instance, the descriptors selected in the Caco-2 permeability model are PEOE Charge BCUT, GCUT logP, molecular refractivity, number of basic atoms, and Van der Waals basic surface area. It makes sense that all of these properties, in particular logP, charges, and molecular refractivity, are highly correlated with the permeability of the compounds. While GCUT logP contributes positively to permeability, the other four have negative impact on permeability. Such relationship is demonstrated by the comparison of quinidine (Caco-2 P = -4.69) and ranitidine (Caco-2 P = -6.31) where quinidine has much higher GCUT logP while the other four properties have lower values than ranitidine. Similar observations have been obtained in the case of hERG and TNBC studies.

Some cheminformatics practitioners contend that machine learning has not fulfilled its promise in predicting biological activity [49]. Poor predictivity is a problem in most models, and there are a variety of possible reasons for this including the incorrect assignment of molecular properties, chance correlation, rough response surfaces, and overtraining [24, 25]. With JavaDL we obtained significant improvement of prediction over other machine-learning models, with most of the prediction errors below one unit, which is usually acceptable in drug design and development. For a few compounds our prediction is not ideal. As shown in **Fig. 6c**, the compound 3-methyladenine (structure shown in **Fig. 7**) is identified as an outlier with the worst prediction in MDA-MB-453 cell lines. To elucidate the reason, we calculate the similarity of 3- methyladenine to other compounds in the training dataset based on their Euclidean distance. We found that the distance to the most similar compound (theobromine) is *SD*_*3-methyladenine*_ *= 1*.*49*. Similarly, we randomly selected two compounds (afatinib and UNC-0638 shown in **Fig. 7**) that were accurately predicted (**Fig. 6c)** and computed their corresponding Euclidean distance to the most similar compounds. We obtained *SD*_*afatinib*_ *= 0*.*68* and *SD*_*UNC-0638*_ *= 0*.*80*, respectively. Using the Tanimoto coefficient (TC), we observed the similar trend: for 3-methyladenine the highest TC = 0.40 (to theobromine); for afatinib and UNC-0638, TC = 0.61 (to pelitinib) and TC = 0.50 (to XL-647), respectively. This indicates that when there are similar compounds in the training set, the prediction is relatively accurate, in agreement with the general principle of QSAR that similar structures tend to have similar activities.

**Figure 7.**
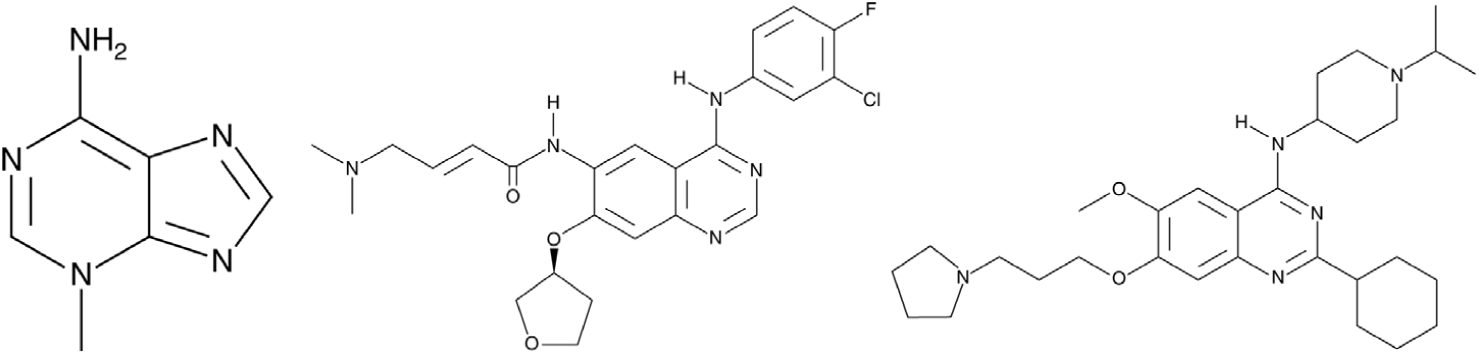
From left to right is 3-Methyladenine, Afatinib and UNC-0638, respectively. 3-Methyladenine was not accurately predicted while the other two compounds obtained accurate prediction of their activity in TNBC cell line.

Our java-based DNN program JavaDL is not limited to drug response predictions; rather it can be very easily used for studies of large genomic and bioinformatics data to predict gene-disease association, identify biomarkers with whole genome profiling, and develop treatment algorithms based on patient response to drugs. Our promising results shown here suggest that JavaDL can be used as a general tool for the discovery and design of biologically active agents as well as for many other types of biomedical research.

## Supporting Information Available

The file for input parameters for training data set HCC_1937 is named “HCC_1937.xml”. The node labeled “net” denotes the configuration of the deep neural networks. It is also available online for download, along with other example files at https://imdlab.mdanderson.org/JavaDL/JavaDL.php.

## Acknowledgements

Special thanks to the Deeplearning4j and CDK toolkit development teams for providing us academic licenses. We also thank the high-performance computing (HPC) resources provided by MD Anderson RIS and Texas Advanced Computing Center (TACC).

